# Differential Gene Expression in MRI-classified Glioblastoma

**DOI:** 10.1101/2024.06.21.600091

**Authors:** Christopher T. Rhodes, Yufeng Wang, Chin-Hsing Annie Lin

## Abstract

Previous characterization of the genome and transcriptome of glioblastoma (GBM) has revealed molecular alterations that potentially drive GBM pathogenesis and heterogeneity ^1-6^. These open-resources are evolving, such as The Cancer Genome Atlas (TCGA) and The Cancer Imaging Atlas (TCIA) at the National Institute of Health comprising a large cohort of molecular and MRI data. Yet, no report deciphers the link between molecular signatures and MRI-classified GBM. The necessity to re-form molecular and imaging data motivated our computational approach to integrate TCIA and TCGA datasets derived from GBM. We uncovered common and distinct molecular signatures across GBM patients and specific to tumor locations, respectively. Despite heterogeneity in GBM, the top 12 genes from our analysis highlights that the dysregulation of a subset of neurotransmitter receptor or transporter and synaptic activity is common across GBM patients. The coherent layer of imaging and molecular information would help us stratify precision neuro-oncology and treatment options in ways that are not possible through MRI or genomic data alone. Our findings provide molecular targets in the disrupted neurocircuit of GBM, suggesting imbalanced excitation and inhibition. Given the fact that GBM patients exhibit similar symptoms resembling patients with neurodegenerative diseases and seizures, our results supported the hypothesis-GBM in the context of neurological disorders beyond a solely cancerous disease.

## INTRODUCTION

High-grade gliomas including glioblastoma (GBM) are devastating primary brain cancer with limited treatment options. The current life-prolonging measures are surgical resection followed by radiation and chemotherapy ^7^ ^8^. Yet, tumor recurrence is possible as tumors can diffusely invade and infiltrate the surrounding brain tissue, making complete surgical resection difficult ^7,9-11^. Mounting collections of molecular analysis had identified several genetic mutations and/or variants in GBM that may predispose an individual to cancer promoting events. Nonetheless, tumor heterogeneity that cells from the same tumor harbor different mutations and exhibit distinct chromatin versus epigenetic status presents challenges to conventional or currently available treatments ^1,12-15^. Within heterogeneity, subtypes of GBM were presumed to arise from glial cells residing in the adult brain, whereas considerable evidence suggests neural stem and/or progenitor cells (NSPCs) as a plausible origin of tumors ^16-25^. Previous study by MRI suggested the intimate relationship between the tumor location of GBM and specific brain regions harboring NSPCs or glial cells ^26^. From the MRI characterized group I GBM, the hypothesis predicts that dysregulation of adult neural stem cells with astrocyte-like morphology in the subventricular zone (SVZ) may give rise to GBM. Similar hypotheses were also implicated to the remaining groups; group II GBM may arise from highly migratory progenitors within the SVZ, group III GBM may come from local progenitors with limited migration, and group IV GBM may be associated with non-SVZ mature glial cells ^26^. Given the awareness that improper control of a fine balance between proliferation and differentiation may contribute to tumorigenesis, our previous studies revealed the abundance of stemness signature in MRI-classified group I and II GBM with distinct clinical outcome that highlights poorly differentiated characteristics in these subtypes of GBM ^24,27-29^. All these compelling evidences support the cell of origin hypothesis. Yet, that does not address the profound phenotypes in GBM patients, such as seizures. GBM in the context of neurological disorders necessitates further investigation. Endeavors in precision neuro-oncology had collected an armamentarium of data housed by TCIA and TCGA. These accessible resources prompted our integrative analysis to re-form image and molecular data. In line with patients’ neurological relevance, our pilot study highlights biological processes and potential candidates associated with aberrant neurocircuit in GBM that provide therapeutic avenues to ameliorate GBM sequalae.

## RESULTS

The purpose of this study was to determine whether there are definitive associations between MRI characterized GBM locations and molecular alterations. We first classified the tumor location of GBM from MRI imaging data of TCIA to group GBM I (SVZ^+^; Cortex^+^), GBM II (SVZ^+^; Cortex^-^), GBM III (SVZ^-^; Cortex^+^), and GBM IV (SVZ^-^; Cortex^-^) ^16-26^. We then integrated the TCIA and TCGA datasets to determine the association between gene signatures and tumor location. One major challenge for our computational approach is the consistency of data format across institutions. To perform integrated analysis in this pilot study, we focus on RNA-Seq datasets from TCGA to streamline and standardize comparisons. As comparing one group of GBM versus all others is a stringent method to find gene signatures, the variance stabilization transformation in DESeq2 normalization process was mainly used to detect differential gene expression in each group of MRI-classified GBM. Heatmaps illustrated that top 500 differentially expressed genes associated with GBM I∼IV. GO analysis revealed that top 500 genes involved in the regulation of proliferation, cell death, and differentiation are commonly affected in GBM while specific sets of genes are altered in particular GBM group (Figure 1). For instance, genes with immune function or xenobiotic metabolism are associated with GBM I (SVZ^+^; Cortex^+^) and GBM II (SVZ^+^; Cortex) compared to GBM III and IV. In addition, MIR31HG, a long non-coding RNA as a host gene for miR-31, was found to be decreased in both GBM I and GBM II (Figure 2). MIR31HG is known to interact with Polycomb repressive proteins to modulate senescence and inhibit proliferation ^30^. Given the fact that c-Myc is involved in etiology of different types of cancer ^31,32^, dysregulation of c-Myc was found in some of GBM cases (Figure 3). c-Myc is known to suppress histone demethylation by interacting with the mammalian homolog of LID protein (Rbp2), which possesses demethylase activity on histone 3 lysine 4 ^33-35^. In addition, c-Myc binds to the promoter of WDR5, a WD40-repeat protein within the Trithorax (Trx) group protein harboring methyltransferase activity to catalyze tri-methylation on histone 3 lysine 4 (H3K4me3)^36^. In the context of chromatin and epigenetic status, c-Myc may activate the transcriptional machinery, at least in part, through inducing Trx-WDR5 methyltransferase and/or inhibition of LID/Rbp2 demethylase. Intriguingly, KDM5D, a histone demethylase specific for histone 3 lysine 4 demethylation was found to be downregulated in all groups of I∼IV GBM cases analyzed. While this integrative analysis deemed usable to validate what are the key factors in hierarchal order to determine epigenetic aberrances as a cause or consequence in GBM, it suggests a shift in chromatin and epigenetic landscapes.

**Figure 1.**
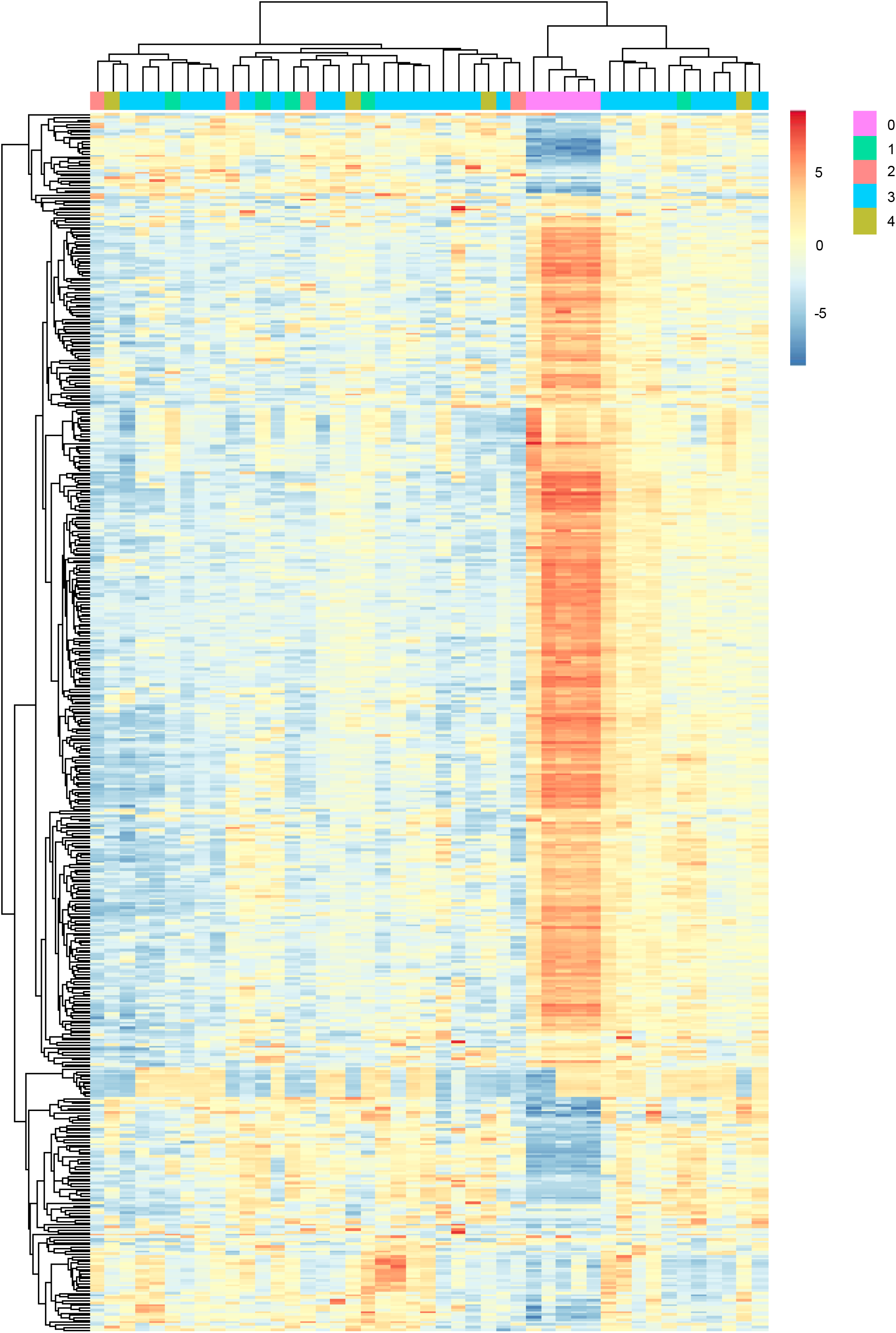
A heatmap represents differential expression of genes in GBM identified from integrated analysis of TCIA and TCGA. Genes used for input are 500 differentially expressed genes with greater than 2-fold change in human GBM corresponding to tumor locations. Inset shows symmetric color scale indicating differences in expression level on log2 scale. Differential expression was analyzed using the function ‘ heatmap.3’ within the R program and unsupervised hierarchical clustering using Euclidian distance metric to identify differences and similarities between genes and between samples (blue shaded versus orange-red shaded clusters in heatmap). Red indicates increased expression of genes in GBM relative to control, blue color indicates decreased expression of genes in GBM compared to control. Dendrogram was determined by hierarchical clustering using Euclidian distance and complete linkage. Color key: 0 normal; 1 GBM I; 2 GBM II; 3 GBM III; 4 GBM IV

**Figure 2.**
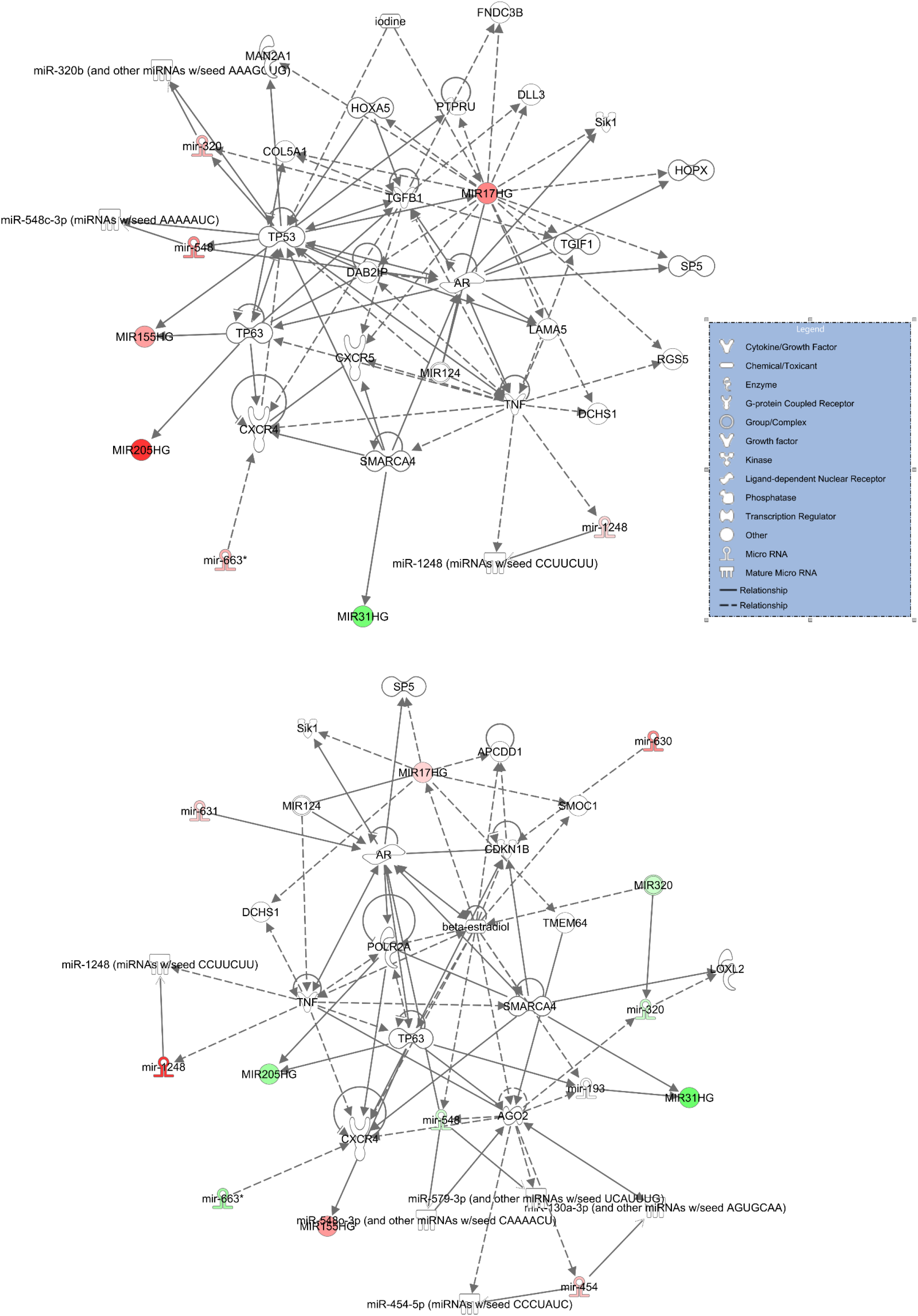
Representative miR network in GBM. Red and green highlight indicates upregulated and downregulated miR in GBM, respectively. Top panel represents miR network associated with GBM tumors contacting both cortex and SVZ while bottom panel represents miR network only in GBM tumors contacting SVZ.

**Figure 3.**
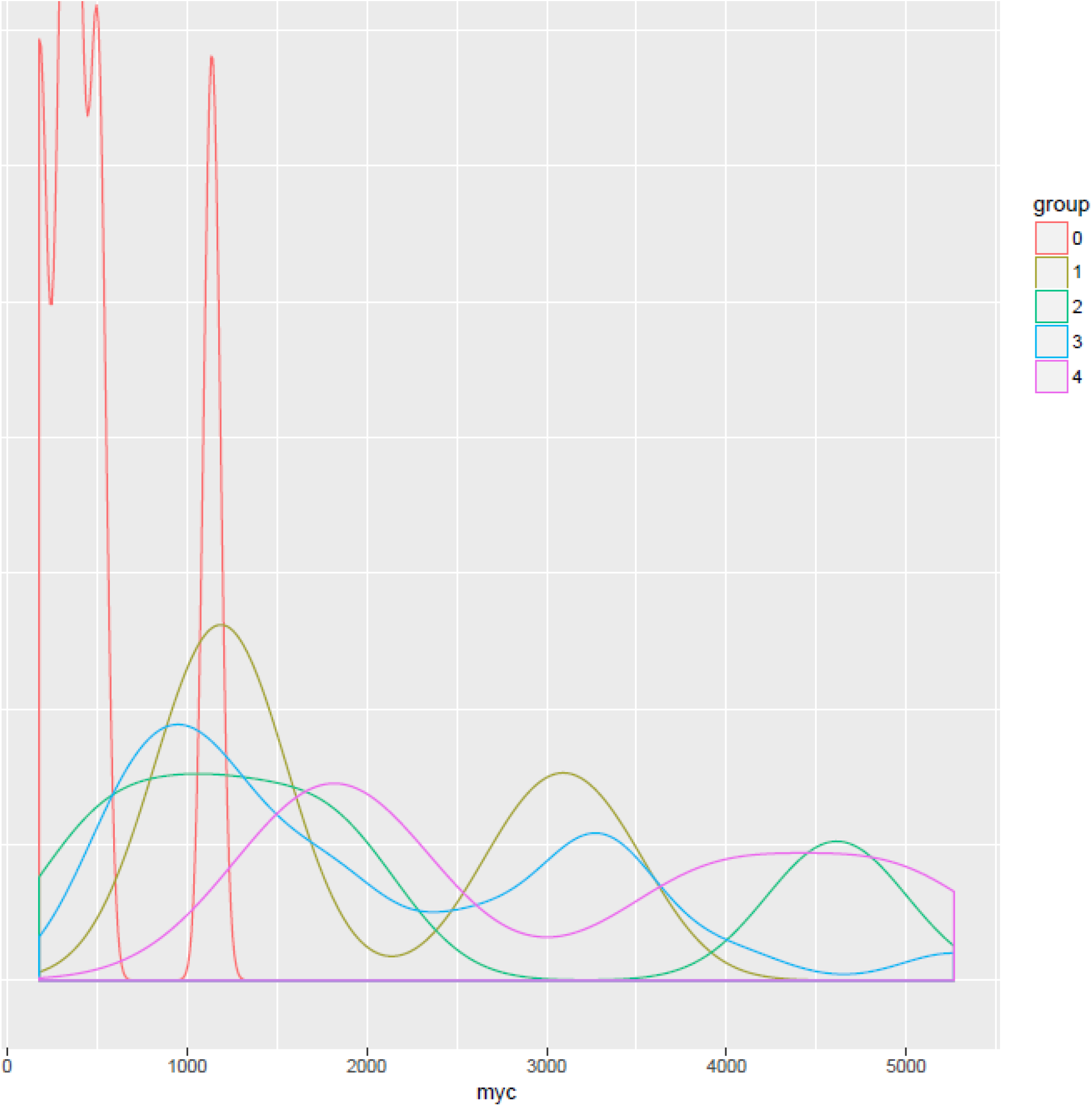
Kernel density estimation (a smoothed version of a histogram) for c-Myc expression per GBM group. X axis represents normalized reads (transcript abundance) of c-Myc, Y axis represents proportion of cases with a given transcript abundance. Group 0 = Controls, Groups 1, 2, 3, 4 = GBM I, II, III, IV based on tumor locations.

Several genes function in transcriptional regulation for nervous system and organismal development were altered across all GBM cases independent of tumor locations. The top network and pathway commonly down-regulated in GBM patients are involved in central nervous system (CNS) differentiation and functional circuit regardless the tumor locations. Among these genes, three are gamma-aminobutyric acid (GABA) receptor subunit receptors (GABRA1, GABRA5, GABRG2). Other CNS-related genes include glutamate transporter (SLC17A7), glutamate receptor (GRIN1, CACNG3), synaptic activity (SV2B, CPLX2), neurofilament polypeptide (NEFL, NEFM), CNS-specific myelin protein (OPALIN), and neuroendocrine secretory protein (CHGA). Therefore, integrative analysis of TCIA and TCGA revealed that a focused set of genes potentially drives neuropathogenesis of GBM. A short list of aberrant overexpression of genes were also identified that include Periostin (POSTN) involved in cell adhesion to promote cell attachment and spreading, iron binding transport protein (LTF), and serum amyloid-A1 protein (SAA1). Whether these commonly dysregulated genes could be potential biomarkers beyond MRI classification that remains to be determined.

## DISCUSSION

Computational approach is not a sole imitability of an existing diagnostic but rather a tool to obtain new information from overwhelming data bank. Hence, given open access repository of brain cancers housed by TCIA and TCGA, we undertook integrative analysis to elucidate the interconnection between tumor locations and molecular signatures. A limitation of this study is variations in the TCIA, in which imaging data were derived from different equipment and reagents used as contrast dyes that resulted in lower numbers of parallel cases of TCGA and TCIA to be analyzed.

The formation of cancerous cells, in some cases, involves somatic mutations of tumor suppressor genes (loss of function) or oncogenes (gain of function). Yet, not all malignant cells contain genomic abnormalities, anomalous epigenetic regulation in cancer cells may be caused by changes in chromatin structure, DNA methylation, histone modifications, and nucleosome remodeling. Because genetic mutations or aberrant gene expression are not always reflected to changes of protein level and do not always predict clinical outcomes, integrating multiple layers of data, like multi-omics or trans-omics would provide a more coherent information. Moving beyond genomic and transcriptomic data, the next integrative analysis would incorporate metabolome and proteomics to expand our complementary assessments. Currently, encompassing multiple institutions had pledged to share targeted proteomic data focus on kinase pathway, while additional comprehensive and quantitative protein measurements would be necessary for multi-omics.

The single-cell qPCR and single-cell RNA-seq for GBM tumors classified in TCGA’ s database showed the presence of multiple subtypes within the tumors. Although informative, it is not surprising from single cell data that multiple molecular pathway-driven-lesions behave distinctly. By contrast, bulk tumor profiling identified the dominant transcriptional program rather than capturing the diversity of transcriptional subtypes within a heterogeneous tumor^15^. It remains invaluable to uncover ubiquitous targets from bulk RNA-Seq; and our approach delineated a list of genes responsible for the disrupted neurocircuit in GBM. Whether neurological abnormalities in GBM are cause and/or consequence during cancer progression remain unclear. Understanding mechanistic underpinnings that alter the functional circuit and governing lesion-induced plasticity in the GBM will advance precision neuro-oncology and provide therapeutic avenues to alleviate GBM sequalae.

## CONCLUSION

Shaping the effort of team science, our audacious intent of integrating tumor imaging and molecular data across GBM cases had revealed a new molecular taxonomy of GBM to advance the improvements in cancer treatment along with mitigating neurological symptom, such as seizures. Among those down-regulated genes, we identified that a set of genes critical for GABAergic system and synaptic activity in GBM. Beyond current antiepileptic medications, our findings offer selective targets responsible for aberrant neurocircuit in GBM that will be of considerable beneficial to those suffering convulsion or seizures. In summary, the work presented here provides emerging insight into coherent health care central to GBM neuropathology, which harbor molecular heterogeneity in conjunction with neurologic complications.

## MATERIALS and METHODS

### Integrated analysis of TCGA and TCIA

#### Data retrieval

To identify GBM cases in TCGA which had corresponding imaging data in TCIA, the National Cancer Institute Genomic Data Commons Data Portal (https://portal.gdc.cancer.gov/) was used to retrieve case IDs for the TCGA-GBM project. Cases IDs were further filtered by selecting only cases that had transcriptome profiling data. Similarly, the TCGA-GBM collection at TCIA was used to search for case IDs which had 1) MRI imaging sequences and 2) case IDs that also appeared in the GDC/TCGA transcriptome profiling database.

#### Classification of GBM tumor location

The MRI imaging data of GBM from TCIA was used to classify tumor locations into four groups. Images were manually reviewed by at least 2 individuals. For each tumor, a T1-weighted sequence was used to classify tumor location. If contrast media (i.e. gadolinium) was used in any T1 sequence, those would be used preferentially over images lacking contrast media. Tumor classification was based on distance of contrast-enhancing lesion (CEL) to nearest ventricle, infiltration of CEL into cortex, volume of CEL, and anatomical location of the CEL. For such, tumor was associated with both SVZ and cortex, only with SVZ, only with cortex, neither SVZ nor cortex, then was assigned numbers GBM I (SVZ^+^; Cortex^+^), GBM II (SVZ^+^; Cortex^-^), GBM III (SVZ^-^; Cortex^+^), and GBM IV (SVZ^-^; Cortex^-^), respectively. This classification (GBM I∼IV) was subjected to integrated analysis with corresponding RNA-Seq data in TCGA. Overall, 142 GBM cases obtained clear MRI were applied for integrated analysis.

#### Gene Expression

To separate the control versus disease and show group specific gene signatures, HTSeq-Counts data for each TCGA-GBM case was downloaded from the NCI GDC portal using the TCGABiolinks package in R/Bioconductor. Sequencing read count matrixes for each GBM group (I, II, III, IV) and control were normalized using Variance Stabilizing Transformation in DESeq2 before differential gene expression analysis which resulted in log2(fold change) values for each group with respect to all other groups. Heatmaps to visualize expression patterns were rendered using pheatmap package (https://cran.r-project.org/package=pheatmap) in R using Euclidean distance and complete linkage.

## GO, Network, and Pathway analysis

DAVID Functional Annotation Tool (DAVID Bioinformatics Resources 6.7, NIAID/NIH) was utilized to perform Gene Ontology (GO) analysis. For GO, a significance cutoff was set at *p*-value < 0.01, including Bonferroni multiple test correction. Functional pathway and network analyses of enriched loci were performed using Ingenuity Pathway Analysis (IPA) (Ingenuity Systems, Redwood City, CA, USA). The top networks and pathways predicted by IPA used cut-off set at *p*-value < 0.01 (calculated by IPA using right-tailed Fisher’ s exact test).

## Authorship statement

CTR performed classification of tumor location and integrated analysis of genomic data. YW performed network and pathway analysis. CAL contributed to approach, material support, project supervision, and manuscript writing. All authors contributed to manuscript preparation.

## ACKNOWLEDGEMENT

We thank Madeleine Moseley for assisting the grouping of tumor location from the TCIA data. We acknowledge the SCORE grant SC3GM112543 from the National Institutes of Health and TRAC award to CAL.

